# Data-driven selection of analysis decisions in single-cell RNA-seq trajectory inference

**DOI:** 10.1101/2023.12.18.572214

**Authors:** Xiaoru Dong, Jack R. Leary, Chuanhao Yang, Maigan A. Brusko, Todd M. Brusko, Rhonda Bacher

## Abstract

Single-cell RNA sequencing (scRNA-seq) experiments have become instrumental in developmental and differentiation studies, enabling the profiling of cells at a single or multiple time-points to uncover subtle variations in expression profiles reflecting underlying biological processes. Benchmarking studies have compared many of the computational methods used to reconstruct cellular dynamics, however researchers still encounter challenges in their analysis due to uncertainties in selecting the most appropriate methods and parameters. Even among universal data processing steps used by trajectory inference methods such as feature selection and dimension reduction, trajectory methods’ performances are highly dataset-specific. To address these challenges, we developed Escort, a framework for evaluating a dataset’s suitability for trajectory inference and quantifying trajectory properties influenced by analysis decisions. Escort navigates single-cell trajectory analysis through data-driven assessments, reducing uncertainty and much of the decision burden associated with trajectory inference. Escort is implemented in an accessible R package and R/Shiny application, providing researchers with the necessary tools to make informed decisions during trajectory analysis and enabling new insights into dynamic biological processes at single-cell resolution.

## INTRODUCTION

Analyses to computationally order related cell types along an underlying dynamic are referred to as trajectory inference or pseudotime analysis(Ji and Ji, 2016). These analytical methods provide users with the capacity to visualize and quantify complex developmental or maturation states using shared gene expression and/or phenotypic protein markers at single cell resolution(Farrell et al., 2018; Kim et al., 2021). While there are numerous trajectory inference methods, obtaining an optimal trajectory remains challenging. A comprehensive review of 45 commonly used trajectory methods, of which there are now more than 100 unique methods at the time of this writing, evaluated each approach’s accuracy of inferred topology, cell ordering, and differential feature expression (Saelens et al., 2019). While no method universally outperformed others, some methods showed distinction in specific metrics. For instance, Monocle (Trapnell et al., 2014) and PAGA (Wolf et al., 2019) best captured the underlying trajectory structure, while Slingshot (Street et al., 2018) most accurately ordered cells. Yet, while this benchmark assists users in narrowing down the top methods for their application, each method requires additional data processing steps and its own set of hyperparameters.

For example, all trajectory methods recommend or require feature selection and dimension reduction prior to trajectory estimation. Feature selection involves subsetting the total number of genes to a smaller set of informative genes, typically the most highly variable genes. However, the decisions of how many highly variable genes and what metric should be used to score gene variability are left to the user. Dimension reduction approaches project the high-dimensional dataset into a low-dimensional (typically two-dimensional) space, with most trajectory methods using either principal component analysis (PCA)(Ji and Ji, 2016), t-distributed stochastic neighbor embedding (t-SNE) (Maaten and Hinton, 2008), uniform manifold approximation and projection (UMAP) (McInnes, Healy and Melville, 2020), or diffusion maps (Haghverdi, Buettner and Theis, 2015). Although, a number of methods, including popular and well-performing ones such as Slingshot(Street et al., 2018), allow users complete flexibility in choosing a dimension reduction technique.

Besides the decision responsibility faced by users in an analysis, the issue lies in the fact that these decisions may lead to significant and unanticipated repercussions (**Fig. 1**). Trajectories inferred using different dimension reduction techniques or numbers of highly variable genes may result in similar biologically reasonable orderings of cell-types clusters along a trajectory (**Fig. 1A**). However, even between visually similar trajectories, the distribution of cells along the trajectory can be highly inconsistent, (**Fig. 1B**) which in downstream analysis, when identifying genes that are dynamic along pseudotime, may lead to contradictory estimates of gene dynamics (**Fig. 1C**). A recently published tool towards this end was designed to facilitate selection of an optimal tree-shaped trajectory by examining cell connectivity(Smolander, Junttila and Elo, 2023). However, it was designed for a specific trajectory inference tool, topology type, and only considers one aspect of the trajectory. Overall, there is little guidance on how various processing choices or hyperparameter settings might affect trajectory estimation, in general, and a lack of quantitative information on whether a particular trajectory is even a good fit to the dataset. As a result, researchers tend to rely on pre-existing knowledge and subjective visual assessment, potentially biasing the analysis and limiting discovery.

**Fig. 1:**
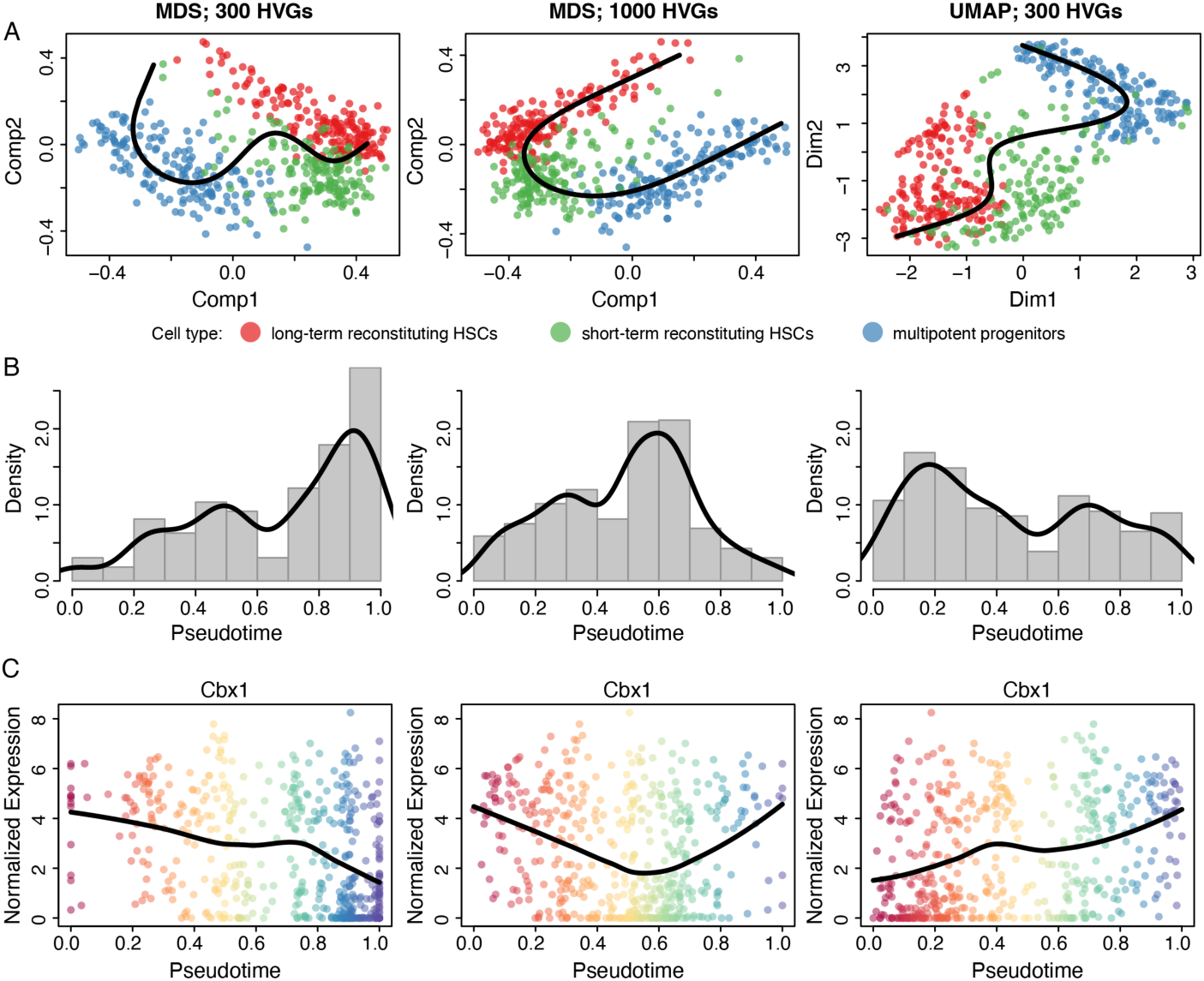
Analysis choices significantly impact trajectory estimation in scRNA-seq data. For various choices of selected genes and dimension reduction methods, trajectory inference and pseudotime estimation was performed on a scRNA-seq dataset of hematopoietic stem cells (Kowalczyk et al., 2015). **A**. Dimension-reduced spaces and estimated trajectories with cells colored by cell type. **B**. Pseudotime distributions for each set of analysis choices. **C**. Normalized gene expression as a function of pseudotime for *Cbx1*. Cells are colored by pseudotime, i.e. their location along the trajectory. Abbreviations: MDS = Multidimensional Scaling, UMAP = Uniform Manifold Approximation and Projection, HVG = Highly Variable Gene.

While benchmarking studies are useful in assessing general method performance and accuracy, evaluation metrics can provide independent assessments of performance when no ground truth exists. For example, the silhouette statistic and stability measures have been utilized to evaluate clustering of cells in scRNA-seq datasets (Zappia and Oshlack, 2018; Duò, Robinson and Soneson, 2020; Yu et al., 2022; Leary et al., 2023). These metrics assess dataset-specific performance in terms of good or desirable properties, e.g., smaller within and larger between cluster distances. While these types of evaluation metrics allow for more-informed decision making when clustering scRNA-seq data, they are not applicable to the continuous nature of trajectory analysis.

To this end, we introduce Escort, a framework for evaluating the impact of choices in trajectory inference and guiding users through the estimation of trajectories in scRNA-seq data. Escort first evaluates a dataset’s overall suitability for trajectory inference and provides dataset-specific summaries allowing users to thoroughly evaluate their analytic approach. Then, based on evaluation metrics we developed to quantify desirable trajectory properties, Escort classifies sets of processing choices as recommended or not-recommended for trajectory estimation. We expect Escort to reduce the decision-making burden in trajectory inference analysis, resulting in less biased and more accurate single-cell trajectories. Escort is available as an R package and available for use as an integrated R/Shiny application, which can be downloaded via Github (https://github.com/xiaorudong/Escort).

## RESULTS

### Trajectory estimation is susceptible to varied analysis choices

In an effort to create a common framework for cell trajectory analysis, we first demonstrated that processing decisions such as choosing different numbers of highly variable genes and different dimension reduction techniques can affect an inferred trajectory. Although these processing steps are only a subset of specific decisions users face, they were chosen for their near universality within trajectory inference methods. We considered eight different simulation scenarios consisting of differences in simulators for scRNA-seq data and trajectory topologies (**Table 1**) (Saelens et al., 2019; Bacher et al., 2022). We refer to each combination of analysis choices as an ‘embedding’ as the vast majority of trajectory estimation methods construct the trajectory in two-dimensional representations. In general, an embedding encompasses all processing choices of interest - including the dimension reduction- and is the dataset on which any particular method estimates the trajectory. In the simulation, the embeddings under evaluation differed in their proportion of highly variable features selected (20%, 40%, 100%) and dimension reduction technique (t-SNE, UMAP, MDS). To avoid over-generalizing, we used both Slingshot(Street et al., 2018) and Monocle3(Cao et al., 2019) to infer trajectories for each embedding. Accuracy of the estimated trajectories was assessed by two different metrics (Kendall rank correlation and mean squared error), each of which we standardized by ranking embedding performance within datasets independently. Overall, there was no clear advantage observed for any specific dimension reduction algorithm, percent of highly variable genes, or combination thereof in terms of performance across datasets (**Fig. 2**). This demonstrates that it is not feasible to simply pre-select a best dimension reduction algorithm or percent of highly variable genes for every analysis or trajectory method.

**Fig. 2.**
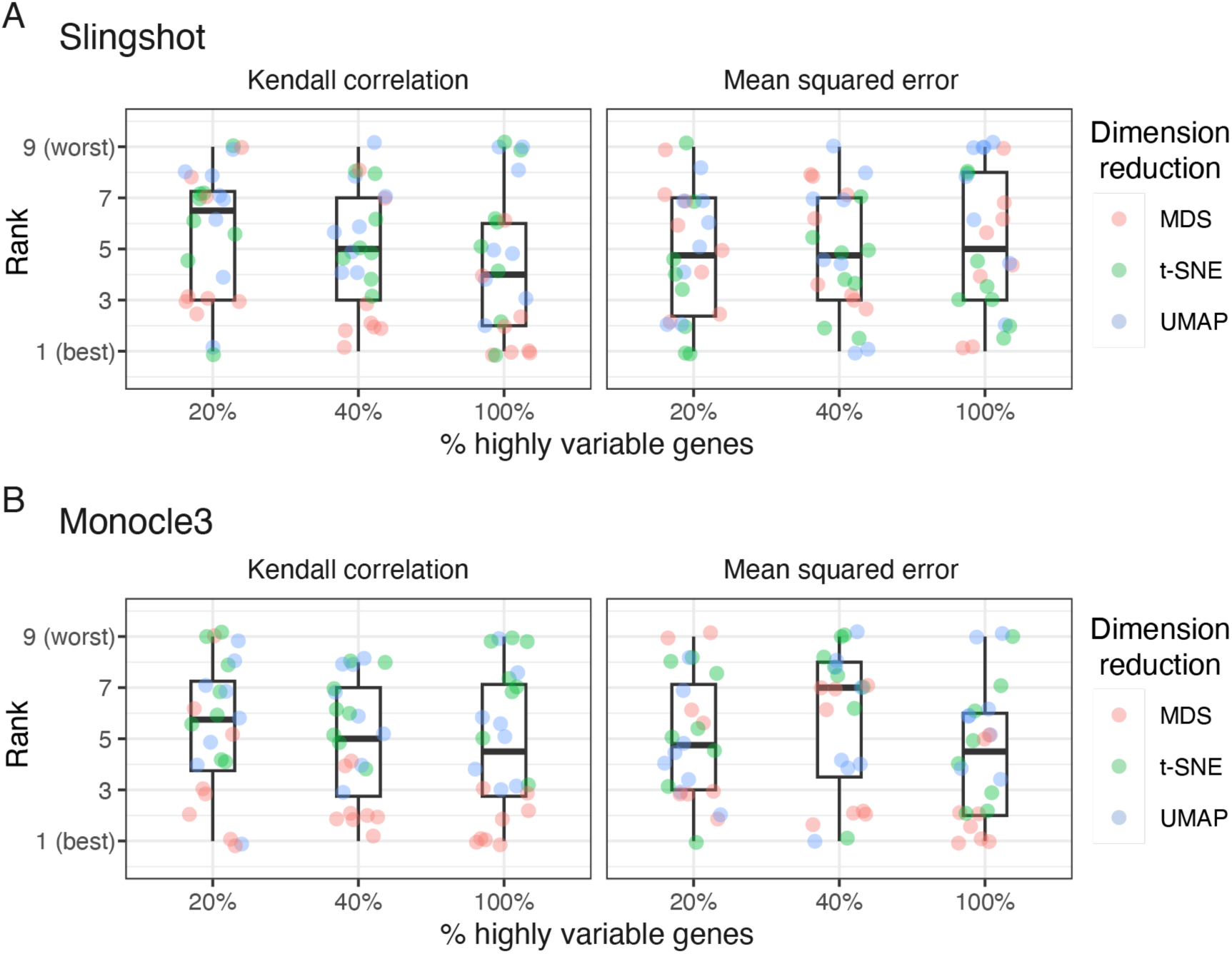
Trajectory accuracy is impacted by different dimension reduction algorithms and inclusions of highly variable genes. **A**. The performance of different embeddings across all eight simulated scenarios is shown. Embeddings were ranked within each dataset separately for the three metrics. The ranks were scaled so that a lower rank indicated better within-dataset performance. **B**. Similar to A using Monocle3.

**Table 1.** A summary of simulated scRNA-seq data datasets.

| Simulation | Method | Trajectory type | Number of cells after filtering |
| --- | --- | --- | --- |
| sim-1 | Scaffold (Bacher et al., 2022) | Linear | 500 |
| sim-2 | dyntoy, available at https://github.com/dynverse/dyntoy | Linear | 263 |
| sim-3 | Splatter (Zappia, Phipson and Oshlack, 2017) | Linear | 174 |
| sim-4 | dyntoy, available at https://github.com/dynverse/dyntoy | Bifurcation/Tree | 748 |
| sim-5 | dyntoy, available at https://github.com/dynverse/dyntoy | Multifurcation/Tree | 1617 |
| sim-6 | dyntoy, available at https://github.com/dynverse/dyntoy | None (Disconnected) | 251 |
| sim-7 | dyntoy, available at https://github.com/dynverse/dyntoy | None (Disconnected) | 674 |
| sim-8 | Scaffold (Bacher et al., 2022) | None (Homogenous) | 500 |

### Framework to identify recommended embeddings for trajectory analysis

Given the dataset-specific performance across trajectory analysis decisions, we developed Escort, a data-driven evaluation framework to guide users through trajectory analysis by providing evaluations of sets of analysis choices (**Fig. 3**). In the first step, Escort assesses whether the data support the existence of a trajectory. Non-computational scientists often struggle with the first step of constructing a trajectory, which is deciding whether fitting a trajectory is appropriate for their dataset. There are two scenarios where trajectory fitting is not well-suited: when cells represent biologically distinct cell types or datasets having insufficient cellular heterogeneity (**Fig. 3A**). Biologically, datasets consisting of distinct cell types indicates that the underlying biological processes are separate or that the experiment did not capture sufficient intermediate-stage cells. Datasets lacking cell heterogeneity may indicate low sensitivity in the experimental assay or excessive technical variability in the dataset. In either case, Escort will flag the dataset in the first step and return a summary with suggestions to the user specifying how to proceed. If a trajectory signal is detected, in the next step the analysis choices (embeddings) that could be used for trajectory estimation are evaluated. Users are able to select a number of default embeddings for consideration or input their own embeddings directly. In this step, Escort quantifies how effectively an embedding preserves relationships between cells in the high-dimensional data and assesses the distribution of points in the two-dimensional cell graph. Based on these two evaluations, an embedding may be classified as non-recommended or passed to the next step (**Methods**). For the third step, embeddings are evaluated in the context of a specific trajectory inference approach in order to allow for consideration of method-specific hyperparameters. Specifically, Escort evaluates the proportion of cells likely to have an ambiguous projection along a trajectory. An overall performance score is calculated based on metrics in the second and third steps, and then the embeddings are classified as recommended or non-recommended. Recommended embeddings are those which are likely to generate trajectories having a better fit to the data and more accurately reflect the underlying dynamic biological processes. Non-recommended embeddings are those that are unlikely to generate accurate trajectories based on the evaluations.

**Fig. 3:**
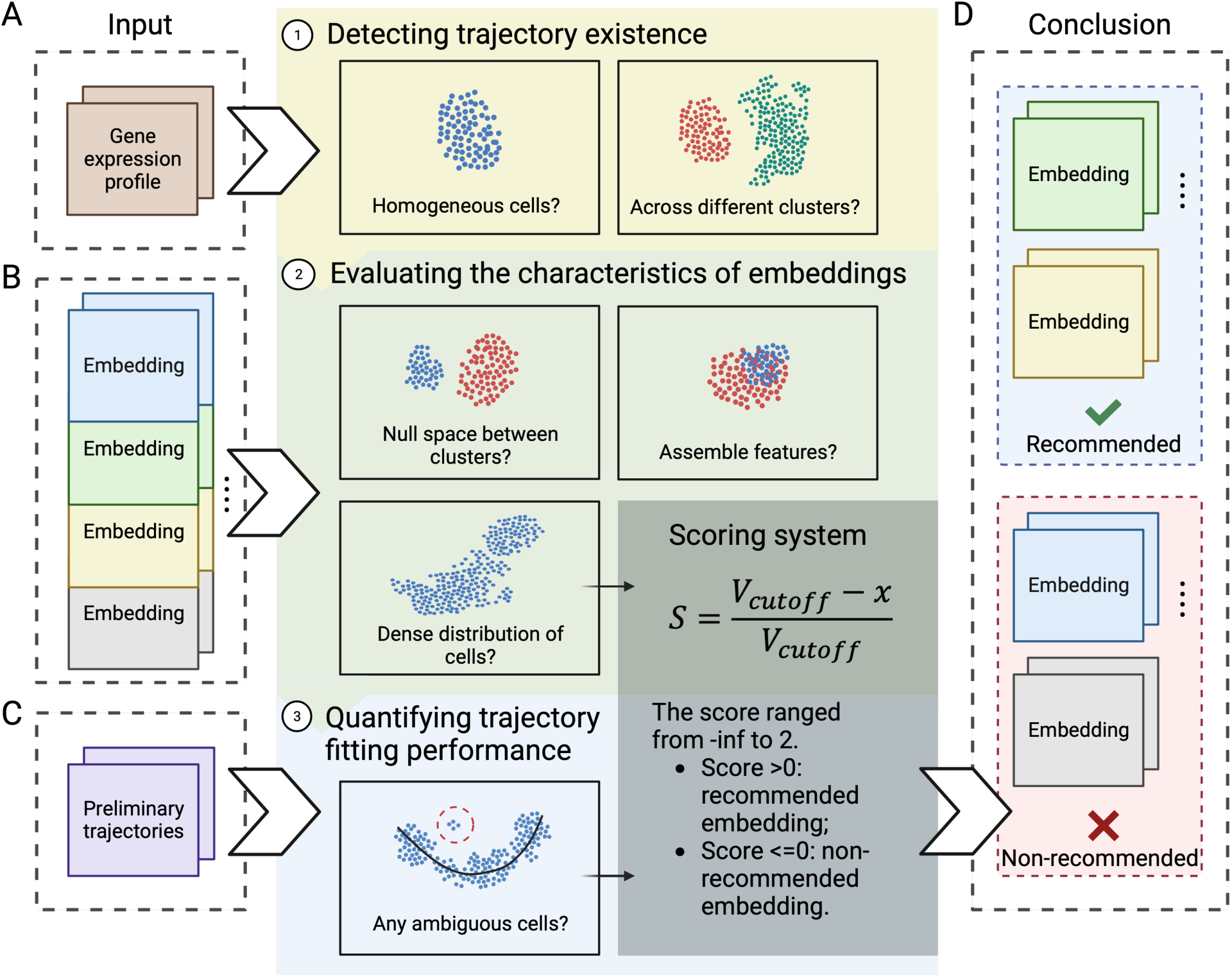
Overview of ESCORT. Schematic of the Escort workflow. **A**. The first step detects the presence of a trajectory signal in the dataset before proceeding to evaluations of embeddings. **B**. Various metrics are using to evaluate user-defined embeddings regardless of the ultimate trajectory inference method to be used. **C**. In the final step, the preferred trajectory inference method of the user is used to fit a preliminary trajectory to allow the evaluation of method-specific hyperparameters. **D**. Based on the overall score, embeddings are classified as either recommended or non-recommended.

### Escort distinguishes embedding quality in simulations

We demonstrate the effectiveness of Escort in evaluating various embeddings on the eight different simulation scenarios (**Table 1**). We used Slingshot to construct trajectories for each embedding, although the results were consistent with Monocle3 (**Supplementary Fig. 1**). Accuracy was assessed in terms of cell order and total order error by comparing the estimated trajectory to the ground truth via Kendall rank correlation and mean squared error. Overall, the recommended embeddings tended to produce more accurate trajectories. Specifically, the recommended embeddings had higher correlation and lower error than non-recommended embeddings (**Fig 4A**). The simulations with no true trajectory were all detected and flagged by Escort in the first step and their embeddings generated significantly less accurate trajectories than even those by the non-recommended embeddings. The Escort score also correlates well with the individual accuracy measures indicating that a higher Escort score reflects higher trajectory accuracy (**Fig 4B**).

**Fig. 4.**
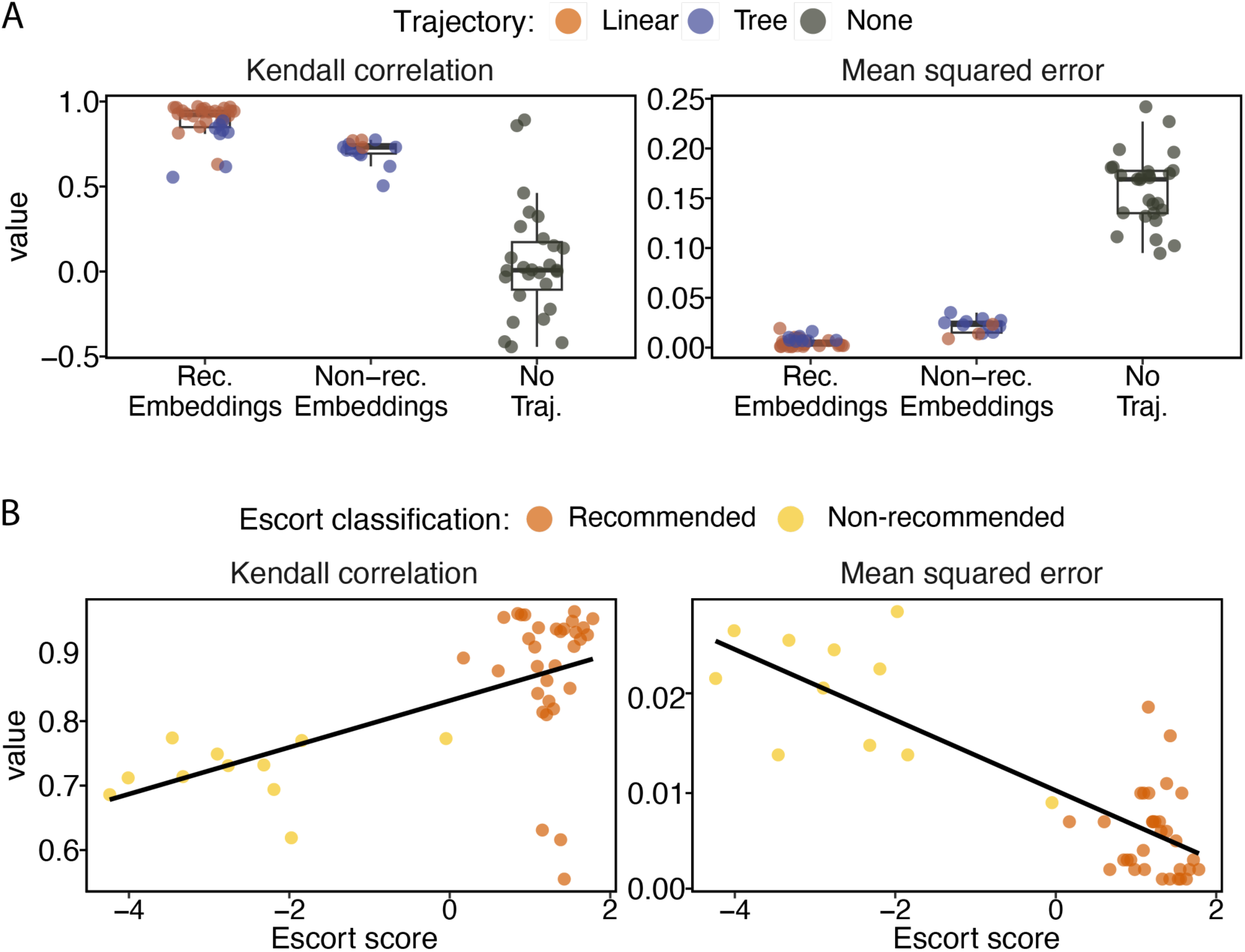
Trajectory assessment performance of Escort on simulated datasets. **A.** The accuracy of trajectories generated on nine different embedding options for each of the eight simulated datasets is shown for different metrics: Kendall rank correlation and mean squared error. Simulated scenarios differ in terms of true trajectory topology (denoted by color) and simulator methods. The y-axis displays the values for the accuracy metric. **B**. Each embedding’s Escort score (x-axis) versus the value for each accuracy metric (y-axis) are shown and colored according to their classification by Escort.

### Escort guides decision-making for trajectory inference

Additionally, we analyzed five scRNA-seq datasets obtained from publicly available sources and encompassed a range of biological contexts (**Table 2**). While the scRNA-seq datasets do not have a ground truth in the sense of a known trajectory, we chose datasets that had some degree of biologically relevant time-ordered samples allowing us to evaluate a trajectory’s fit to the data. Despite the increased noise and complex data structure, Escort is still able to distinguish embedding quality (**Fig. 5 and Supplementary Fig. 2**). The recommended embeddings have a significantly larger accuracy compared to the non-recommended embeddings. The accuracy measures are nosier for these datasets compared to the simulations due to the small number of true times in each, however higher accuracy is still correlated with a higher Escort score (**Fig. 5B**).

**Fig. 5.**
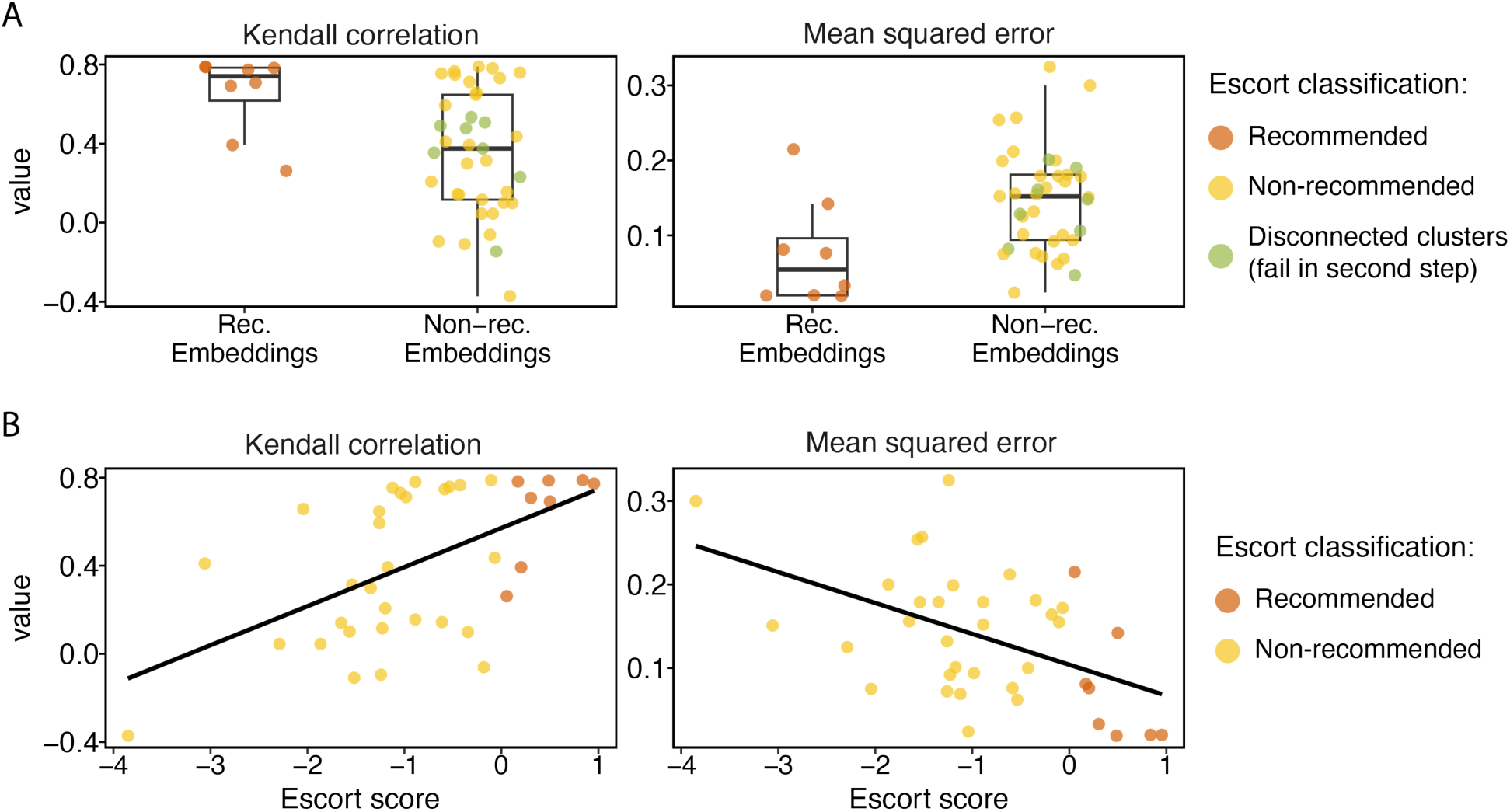
Trajectory assessment performance of Escort on public datasets. **A.** The accuracy of trajectories generated on nine different embedding options is shown for five publicly available datasets assessed using different metrics: Kendall rank correlation and mean squared error. The colors distinguish each embedding classification by Escort, in addition to those embeddings that failed in the second step. The y-axis displays the values for accuracy metrics. The x-axis corresponds to recommendations generated by Escort. **B**. Similar to A with the x-axis showing the Escort score.

**Table 2.** A summary of public scRNA-seq datasets. Datasets sc-2 through sc-5 were downloaded from Saelens et al. (Cannoodt et al., 2018b)

| Dataset | Organism | Accession | Number of cells after filtering | Number of time points |
| --- | --- | --- | --- | --- |
| sc-1 | Human | E-MTAB-3929<br>(Petropoulos et al., 2016) | 1469 | 5 |
| sc-2 | Mouse | GSE59114<br>(Kowalczyk et al., 2015) | 492 | 3 |
| sc-3 | Mouse | GSE99951 (Sloan et al., 2017) | 448 | 4 |
| sc-4 | Mouse | GSE87375 (Qiu et al., 2017) | 321 | 6 |
| sc-5 | Human | GSE86146 (Li et al., 2017) | 647 | 12 |

We also found that that the scRNA-seq datasets tended to have more non-recommended embeddings compared to the simulated datasets. Non-recommend embeddings are a result of having a negative Escort score, and additionally, when the embedding fails to sufficiently preserve complex cell relationships, it is immediately classified as non-recommended. For example, we found that UMAP frequently generated embeddings that exhibited disconnected clusters or grouped small clusters together in unexpected ways (Charrout, Reinders and Mahfouz, 2020). These more complex datasets also have more irregularly spaced sampling times, for example, the sc-3 dataset (**Table 2**) had single-cells measured at days 100, 130, 175, and then day 450. If too few cells are represented from intermediate progression sampling times or cell states, it is unlikely any embedding can overcome experimental design limitations and consistently generate robust connected trajectories.

### Escort guided trajectory analysis of hypertrophic chondrocyte transdifferentiation

Hypertrophic chondrocytes were once thought to be the terminal state of chondrocyte fate prior to apoptosis, yet recent research suggests these cells undergo transdifferentiation towards an osteoblast-like state or other marrow associated cells (Sa, W and N, 2021). To better understand the fates of hypertrophic chondrocytes, Long et al. (Long et al., 2022) utilized lineage-tracing mouse-reporter models to isolate *Col10a1*-expressing cells and their descendants. The authors performed trajectory inference on their scRNA-seq data using default settings in Monocle3, resulting in a singular lineage linking hypertrophic chondrocytes to osteoblasts. Using the original paper’s publicly available dataset, we sought to examine whether Escort would recommend a more optimal embedding that might provide additional insight into this differentiation process.

Prior to trajectory estimation, we found that one of the initial cell clusters (cluster 7) appeared to be low quality cells with very low sequencing depth (**Suppl. Fig. 3**). Removing these cells from the dataset resulted in a highly similar, single-lineage trajectory using the default settings in Monocle3 (**Fig. 6A**). However, for our analysis, given the interest in the multi-fate hypothesis of hypertrophic chondrocytes, we opted to use Slingshot to fit trajectories on both the original embedding and an Escort recommended embedding (**Fig. 6B&C**). Escort evaluated 13 various embedding options. The original embedding scored sub-optimally and was ranked in the bottom quartile among all embeddings (**Supplementary Table 1**). An Escort-based trajectory was fit using Slingshot with the highest-scoring embedding option.

**Fig. 6.**
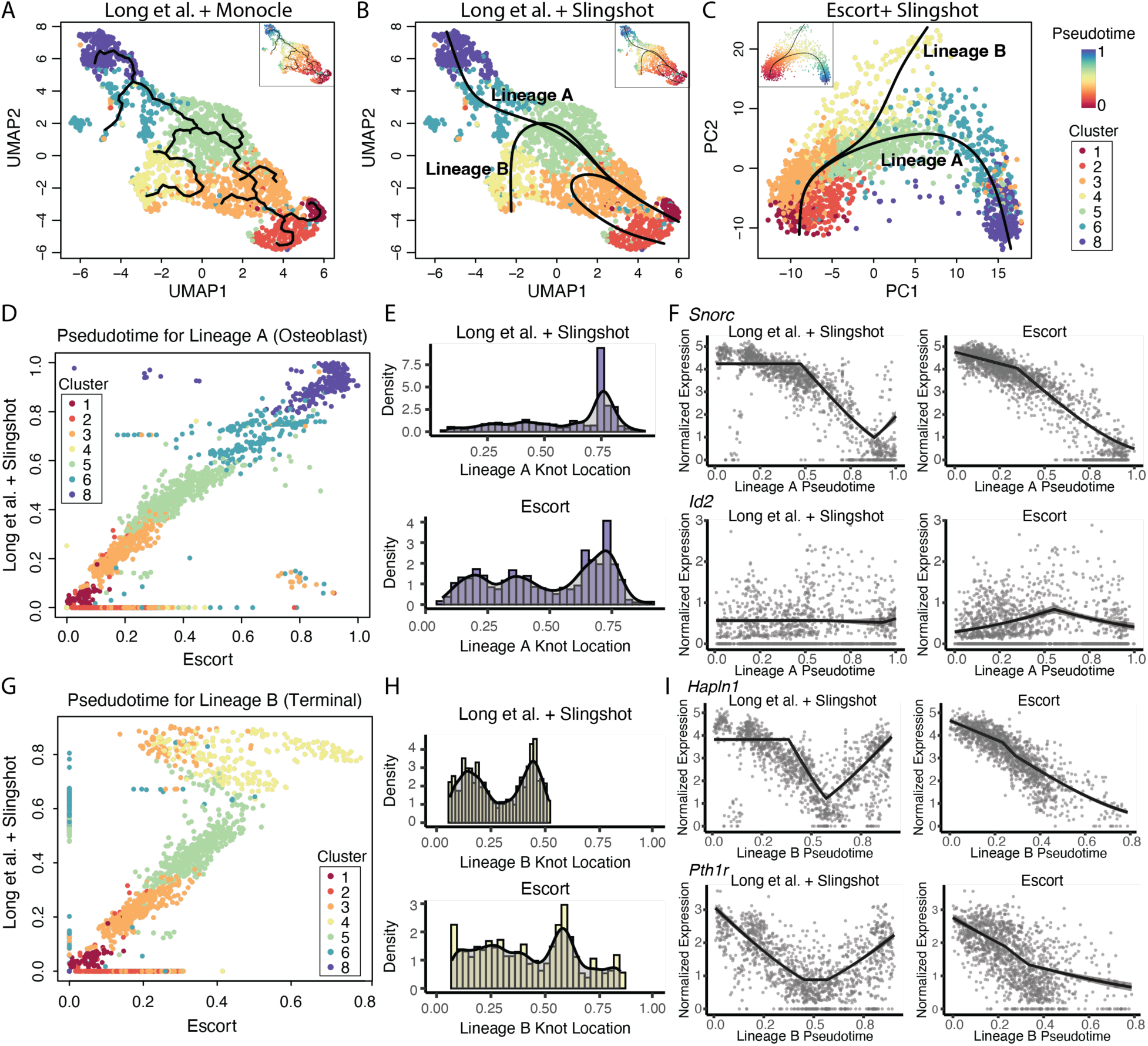
Analysis of transdifferentiation of hypertrophic chondroblasts using an Escort guided trajectory. **A.** UMAP of the original paper’s embedding and the Monocle3 based trajectory. **B.** UMAP of the original paper’s embedding using Slingshot to fit a trajectory. **C.** Escort recommended embedding using Slingshot to fit a trajectory. **D.** Correlation of pseudotime between the two Lineage A trajectories. **E.** Distribution of knots across all significantly dynamic genes for Lineage A. **F.** Gene expression as a function of pseudotime for *Snorc* and *Id2.* **G-I**. Similar to **D-F**, but for Lineage B.

Topologically, the two trajectories were highly similar, although the original embedding generated three lineages while Escort’s embedding generated two (**Fig. 6B-C**). Based on the clusters along each branch, we labeled the two shared lineages as Lineage A and Lineage B. Based on the original author’s annotations, Lineage A follows hypertrophic chondrocytes to osteoblasts and Lineage B links hypertrophic chondrocytes to terminal hypertrophic chondrocytes. The third lineage, only observed using the original embedding, was enriched for VEGFA signaling and protein transport, which indicated heterogeneity within the dedifferentiating hypertrophic chondrocytes (**Supplemental Data 1**). For our analysis, we focused on comparing how the two embedding options differed in downstream analysis when characterizing gene dynamics in Lineages A and B. For each lineage, we applied scLANE to identify differentially dynamic genes(Leary, J. and Bacher, R., 2023). scLANE models each gene’s expression as a function of pseudotime using a modified multivariate adaptive regression spline approach. The scLANE model identifies locations of significant expression changes along pseudotime (referred to as knots) and estimates the expression slopes along each interval.

For Lineage A, the osteoblastic fate, the two trajectories had similar overall topology and pseudotime distributions, however, the Escort-based trajectory resulted in 12% more genes identified as significantly dynamic compared to the original trajectory (**Fig. 6D**). The distribution of knots, indicating locations of major expression change along the lineage, was multi-modal for the Escort trajectory whereas the original trajectory had a unimodal distribution with a high concentration of knots when transitioning between clusters 6 and 8, indicating disconnectedness. Although the majority (> 85%) of shared differentially dynamic genes had similar trends, the Escort-based dynamics were less noisy and captured more subtle expression changes as seen in chondrogenic differentiation genes *Snorc* (Heinonen et al., 2011) and *Id2* (Sakata-Goto et al., 2012) (**Fig. 6F**).

The lineage towards more terminal hypertrophic chondroblasts, Lineage B, varied substantially between the two trajectories (**Fig. 6G**). The original trajectory included cells from Cluster 6, thought to be a transitory stem and progenitor-like cell population, which were predominantly only in Lineage A in the Escort trajectory. The knot distribution for the original trajectory had a large proportion of knots appearing in the initial transition from clusters 1 to 3, likely due to a small gap in the pseudotime, and a large proportion of knots appeared at the transition between clusters 5 to 6 (**Fig. 6H**). These differences in pseudotime distributions lead to 35% of all shared significantly dynamic genes having opposing dynamic trends (upregulation versus downregulation). Genes upregulated in the original trajectory, but downregulated in the Escort trajectory were enriched for cartilage and skeletal system development. Genes upregulated in the Escort trajectory and downregulated in the original trajectory were enriched for regulation of RNA splicing and the EGFR1 pathway, which stimulates terminal differentiation and apoptosis in chondrocytes (X et al., 2013) (**Supplemental Data 2**). Lineage B in the Escort trajectory clearly represented cells heading towards the apoptotic fate, whereas the original trajectory contained blended, less distinct signals among its lineages. The Escort-guided analysis led to a trajectory that more accurately characterized the underlying transdifferentiation of hypertrophic chondrocytes, with two clearly distinct lineages towards osteoblastic fate and terminal apoptosis, whereas the embedding used in the original analysis was only able to characterize the osteoblastic linage.

## DISCUSSION

While all-inclusive or automated analysis pipelines are desirable for ease of use, recent discussions have highlighted issues regarding careful inference in single-cell data (Johnson, Kath and Mani, 2022; Neufeld et al., 2022; Chari and Pachter, 2022). In this context, it is crucial to consider the unique characteristics of the data when selecting among trajectory inference methods and performing processing steps. To address this challenge, we developed Escort to evaluate the impact of choices in trajectory inference while guiding users through the estimation of trajectories in scRNA-seq data analysis. Our framework is implemented in an R package to allow users to evaluate as many embedding options as desired, as well as through an R/Shiny application to guide users more directly through assessing data-specific properties to consider when performing trajectory inference.

Given that trajectory inference methods often differ in their use of dimension reduction techniques, our results observing highly dataset-specific performance of is consistent with that observed comparing methods in the benchmarking report by Saelens et al. (Saelens et al., 2019). In fact, the processing choices for dimension reduction and inclusion of highly variable genes impacted the trajectory accuracy more than the difference between Slingshot or Monocle3 for estimating the trajectory and pseudotime. In addition to standard processing steps, quality control steps may also significantly impact Escort’s embedding evaluations. For example, normalization is a key upstream pre-processing step that adjusts for variations among cells due to sequencing depth. Inadequate normalization may compromise accurate trajectory signal detection (Lun, 2018; Tian et al., 2019). Batch effects also have the potential to introduce technical variability between samples (Leek et al., 2010). Batch effects in scRNA-seq data can result in incorrect trajectory signal assessment and the emergence of batch-specific clusters or trajectories that do not accurately represent biological characteristics (Büttner et al., 2019; Hie, Bryson and Berger, 2019; Lakkis et al., 2021). Thus, we highly suggest that these decisions also be defined explicitly in the embedding to allow for evaluation within the Escort framework.

Additionally, we and others have found that the performance of UMAP is heavily dependent on hyperparameters, specifically the minimum distance between points in the lower-dimensional space and the number of approximate nearest neighbors used for constructing the initial high-dimensional graph (Wang et al., 2021; Xia, Lee and Li, 2023). The choice of these hyperparameters has a significant impact on how UMAP behaves (Liu, 2020; Ehiro, 2023). Hyperparameters such as these can also be incorporated within the Escort evaluation framework.

Finally, Escort is not guaranteed to identify a recommended embedding among the choices being evaluated. Maintaining connected relationships within an embedding is a general challenge and has been previously reported by others (Charrout et al., 2020; Fischer, Burkholz and Vreeken, 2023; Xu et al., 2023). Insufficient experimental designs may also present situations with small discontinuities. Given that the Escort score was highly correlated with accuracy of the trajectory, then when a reasonable number of embedding options have been explored, Escort’s score can be utilized in cases where it may be necessary to choose the best possible option, understanding that there is some unavoidable discontinuity present. In any case, Escort provides users with the ability to identify more optimal analysis choices in the context of trajectory inference.

## METHODS

Our three-step framework aims to improve the accuracy of trajectory estimation guiding users through the decisions involved in fitting a trajectory.

### Step 1: Detecting trajectory existence

The first step of trajectory analysis is deciding whether fitting a trajectory is appropriate for a given dataset. There are two scenarios where trajectory fitting is not appropriate: when cells represent diverse cell types or when cells appear homogeneous.

To identify distinct cell types, we use two single-cell specific approaches to identify clusters: scLCA (Cheng et al., 2019) and SC3 (Kiselev et al., 2017). The scLCA approach estimates the optimal number of clusters and cell clusters in the dataset based on cosine-similarity and spectral clustering (Yu et al., 2022). SC3 is a consensus cluster approach; a consensus matrix is constructed from clustering with multiple distance metrics and then split into a user-specified number of clusters. In scenarios where cell type information is lacking, Escort depends on the optimal number of clusters and their assignments determined by scLCA. Conversely, if prior information regarding cell types is known, Escort utilizes SC3 to compute the cell clusters. Next, Escort computes distances between the cell clusters in the high-dimensional space to evaluate their connectivity. This assessment assumes that large distances between clusters indicates disconnectedness between the cell types. To evaluate connectivity between clusters, let *N* be the total number of cells in the dataset, denoted as *c*_1_, *c*_2_, …, *c_N_*. The Manhattan distance matrix *D_N×N_* is defined as

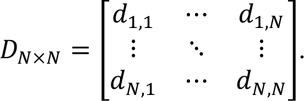

where *d_i,j_* represents the Manhattan distance between cell *c_i_* and *c_j_*.

The *m* cell clusters are denoted as *T*_1_, *T*_2_, …, *T_m_*, such that the cluster *T_k_* consists of a subset of *n*_k_ cells. If *c_i_* ∈ *T_k_*, then 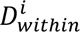 is the set of “within-cluster distances” for *c_i_* such that 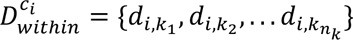 and 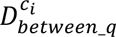 represents a set of “between-cluster distances” for *c_i_* between cells in *T_q_*, given by 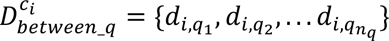, where *q* ∈ {1, 2, …, m} and *q* ∉ *k*. Jaccard index scores are then computed for *c_i_* based on the distributions of 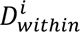 and 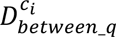, where *q* ∈ {1, 2, …, *m*} and *q* ∉ *k*. Cells with high Jaccard indices across multiple clusters indicate low cluster specificity and their potential presence at the boundaries of clusters. The Jaccard index is recalculated for the top 20% of cells with the highest Jaccard index score within each cluster. Two clusters are considered to be connected if, in pairwise comparisons, there are more than ten cells meeting the 0.3 cutoff criteria for each cluster. We employ the threshold of 0.3 and consider a minimum of 10 cells based on the findings of a simulation study. Otherwise, clusters are considered disconnected.

To assess the homogeneity of the cells in the dataset, Escort analyzes the correlation between the first principal component of the normalized expression data (PC1) and a random subset of the top 100 highly variable genes. This assumes that if a true trajectory signal exists, then it would be encoded within multiple highly variable genes and also captured by the first principal component. A significant correlation between PC1 and more than 46% of the highly variable genes at a significance level of 0.05 was used as our criterion, determined through a simulation study, for identifying a trajectory signal. Significance is assessed via a permutation test with 20,000 iterations of the Spearman correlations accompanied by a false discovery rate correction.

### Step 2: Evaluating the characteristics of embeddings

Once Escort has determined the suitability of trajectory analysis, the second step is designed to evaluate embeddings for performing trajectory inference. Since all methods employ some form of dimension reduction, we refer to the collective set of choices as an ‘embedding’ of the dataset. However, in practice, an embedding may also consist of methods for normalization, selection of highly variable genes, or other method-specific hyperparameters.

The first evaluation of an embedding is the retention of inter-cellular relationships that are present in the high-dimensional data. While we do not expect a (typically) lower dimension embedding to perfectly preserve high dimensional relationships, Escort penalizes severe distortions as the accuracy of trajectory prediction is heavily dependent on the extent to which these relationships are preserved (Saelens et al., 2019). The distance-based method introduced in Step 1 is utilized again to evaluate cell connectivity on the embedding. Since disconnected clusters were resolved in Step 1, a reliable embedding should not exhibit newly disconnected clusters. Consequently, any embeddings found to be disconnected are classified immediately as not recommended for trajectory inference.

For any remaining embeddings, the preservation of similarity relationships in the low-dimensional embedding is evaluating by employing what we define as the “same group level” method. For each cell in the embedding, the three closest cells based on Euclidean distance in low-dimensional embeddings are identified. A cell has a high “same group level” when at least two of its closest neighbors belong to its same cluster as defined in Step 1. The clustering structure observed in the higher-dimensional space is considered to be well-preserved in a given embedding when a large number of cells exhibit a high “same group level”. For example, consider the detection of the three closest neighbors for a cell *c_i_*. The set *NS_i_* = {*c_i_*_1_, *c_i_*_2_, *c_i_*_3_} contains the three closest neighbors for *c_i_*. If *c_i_* ∈ *T_k_* and {*c_i_*_1_, *c_i_*_2_, *c_i_*_3_} ∉ *T_k_*, then the “same group level” is considered 0. If {*c_i_*, *c_i_*_1_} ∈ *T_k_* and {*c_i_*_2_, *c_i_*_3_} ∉ *T_k_*, the “same group level” is considered 1. If the “same group level” is equal to or greater than 2, we conclude that the cell exhibits a high level of similarity with its neighboring cells. The preservation score for similarity relationships is determined by the percentage of cells exhibiting a high level of similarity.

The next embedding evaluation considers cell density. If cells are more uniformly distributed in the two-dimensional embedding space, the trajectory inference method faces challenges in identifying a robust trajectory. Conversely, if cells are denser and exhibit a distinct topology, the trajectory inference method is more likely to generate a well-defined curve. Since methods frequently fit trajectories on two-dimensional representations, we quantify the cell density by calculating “cell coverage area” using the area of the α-convex hull (Pateiro-López and Rodríguez-Casal, 2010). The proportion of this area to a minimum circle enclosing all cells in the two-dimensional embedding space is then calculated.

### Step 3: Quantifying trajectory fitting performance

So far Escort has provided independent evaluations of embeddings for trajectory inference based on general properties, specific methods may impose additional specific graph structure. Thus, the final step for Escort accounts for any method-specific variations in the performance of embeddings.

For each embedding, a rough trajectory is fit using an assumed method. We use Slingshot as the default, however, other methods are easily incorporated at this stage and described in the user vignette. With the embedding-specific trajectory, Escort estimates the proportion of cells positioned along the trajectory such that their projection is ambiguous. For example, trajectories in a U-shape tend to be less accurate due to the presence of cells that map with similar probability to the beginning or end of the trajectory. Each cell’s pseudotime is computed by each trajectory inference method by projecting cells onto the trajectory. Escort calculates each cell’s pseudotime standard deviation based on the closest 10% of its nearest neighbor. Ambiguous cells are then defined as those with an extremely large standard deviation exceeding the upper fence identified by a skew-adjusted approach (Hubert and Vandervieren, 2008). If multiple lineages are present, then this step is performed per lineage, followed by the calculation of the total number of uniquely ambiguous cells.

### Escort scoring

A comprehensive score incorporating the assessments above is used to evaluate the overall performance of each embedding. The score consists of three components: proportion of cells having a high same-group level (step 2), cell coverage area (step 2), and the proportion of ambiguous cells (step 3). Embeddings deemed non-recommended based on the disconnectedness evaluation in the second step are not included in the score comparison.

To establish a realistic and standardized benchmark scoring system, we conducted simulations using Scaffold based on a human pancreas single-cell RNA-seq dataset (Baron et al., 2016; Bacher et al., 2022). We simulated a total of 50 scRNA-seq datasets, each comprising 500 cells, 15894 genes, and including 20% dynamic genes. For each of the simulated datasets, we produced nine different embeddings by employing MDS, UMAP, and t-SNE based on a subset of 2,000, 4,000, or 10,000 highly variable genes. Subsequently, we assessed all generated embeddings by computing both mean squared error (MSE) and Kendall rank correlation coefficients. We identified good embeddings as those with correlation coefficients exceeding 0.85 and MSE less than 0.01. We then fit a beta distribution to the cell coverage area values generated by the good embeddings, as well as a gamma distribution to the proportion of ambiguous cells within these embeddings. We established suitable cutoffs as the 99^th^ percentile of the standard fitted distributions. This led to a cutoff for cell coverage area of 0.53 and for the proportion of ambiguous cells as 0.029. In practice, these cutoffs have proven reasonable.

To classify embeddings as recommended or not recommended based on the score, Escort scales the scores separately for the density and ambiguous components as:

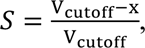

where *x* is the value of cell density or proportion of ambiguous cells. For cell density, we set the *V_cutoff_* to 0.53, while for the proportion of ambiguous cells, the *V_cutoff_* is 0.029. The percentage of cells with a high “same group level” is included in the score as a weight. The total score for each embedding is calculated as

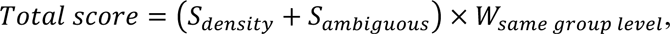

where *S_density_* is the scaled score for the coverage area components, *S_ambiguous_* denotes the scaled score for ambiguous cells component, and *W_same group level_* represents the percentage of cells with a high “same group level.” The score ranges from negative infinity to two, with higher scores indicating better performance. Escort reports embeddings with a score greater than zero as recommended, while those with a score less than or equal to zero as non-recommended. Through this scoring system, users can readily identify more optimal embedding choices to construct a trajectory.

### Application of Escort to simulation and single-cell RNA-seq datasets

We generated two simulated datasets using the Scaffold v0.2.0 R package(Bacher et al., 2022), one consisting of homogenous cells and the other featuring a linear trajectory. The reference data for simulation was obtained from human pancreas single-cell RNA-seq (Baron et al., 2016). The estimateScaffoldParameters() function was utilized to configure all simulation parameters. In the dataset with homogenous cells, no dynamically expressed genes, which are genes influencing cell development, were identified. For the linear structure dataset, we specified that 20% of the genes should be dynamic. The simulateScaffold() function was applied to run the simulations based on the parameters from the estimateScaffoldParameters() function. Additionally, we utilized six simulated datasets provided by Saelens et al., available on Zenodo (https://doi.org/10.5281/zenodo.1443566) (Cannoodt et al., 2018a).

For each simulated dataset, we initially eliminated cells with duplicated ground truth time. Then, we preprocessed each dataset by removing low-quality cells based on extreme outliers (low or high) counts of unique genes, total molecule counts per cell, and a high percentage of mitochondrial genome reads using Seurat v5.0.1(Hao et al., 2023). Genes having less than 3 total counts (or less than 10 for the sc-1 dataset) were also filtered out. The normalization method ‘LogNormalize’ was then applied using the NormalizeData() function in Seurat.

Highly variable genes were identified using scran v1.28.2 (Lun, McCarthy and Marioni, 2016). The modelGeneVar() function identified genes demonstrating higher variability than expected given their mean expression. We selected features based on biological variability by utilizing the “bio” argument in the getTopHVGs() function, picking the top 20%, 40%, and 100% proportion of genes. Then, t-SNE, UMAP, MDS were applied as dimension reduction technique to generate embeddings. t-SNE was performed by Rtsne v0.16 package, involving an initial PCA on the normalized data with the default setting retaining the top 50 dimensions. UMAP was implemented using the R package umap v0.2.10.0 directly on normalized data with default settings. SCORPIUS v1.0.9 (Cannoodt et al., 2016) was used to implement MDS with the spearman distance metric. A trajectory was inferred for each embedding using both Slingshot v2.8.0 (Street et al., 2018) and Monocle3 v1.3.4 (Cao et al., 2019). The mclust v6.0.1 (Scrucca et al., 2023) was used to generate clusters based on hierarchical clustering, and these clusters were subsequently employed as input for Slingshot.

### Analysis of hypertrophic chondrocytes

The pre-processed scRNA-seq data from Long et al., 2022 was downloaded from GSE190616. Escort was applied to the normalized expression matrix for 13 different embedding comparisons (Supp. Table 1), the embedding option using PCA with 10% of the most highly variable genes performed best was selected for further trajectory analysis. Slingshot v2.8.0(Street et al., 2018) was used to fit the trajectory under the Escort embedding option, as well as the embedding used in the original paper’s analysis. The R package scLANE v0.7.8(Leary, J. and Bacher, R., 2023) was used to test each gene’s expression for trajectory differential expression using the default settings. Genes were considered significantly dynamic if they have an overall false discovery rate adjusted p-value < 0.01. Genes were further filtered for enrichment if they had the last segment slope > 5 or < −5. Enrichment was carried out using both Enrichr(Xie et al., 2021), specifically using the BioPlanet 2019 collection and GSEA Molecular signatures database (MSigDB) for mouse(Liberzon et al., 2011; Castanza et al., 2023), specifically the GO biological processes collection.

## Data availability

The processed simulated, scRNA-seq, and hypertrophic chondrocyte datasets are available on Github (https://github.com/xiaorudong/Escort-paper).

## Code availability

Code used to analyze the hypertrophic chondrocyte data along with analysis results is provided on Github (https://github.com/xiaorudong/Escort-paper). The Escort package and tutorials are available on our Github repository (https://github.com/xiaorudong/Escort).

## Acknowledgments

This work is supported by NIH grant R35GM146895 to R.B. and NIH grant P01AI042288 to T.M.B.

## Notes

### Competing Interest Statement

The authors have declared no competing interest.

