## Supplemental Figures and Tables for "Data-driven selection of analysis decisions in single-cell RNA-seq trajectory inference"

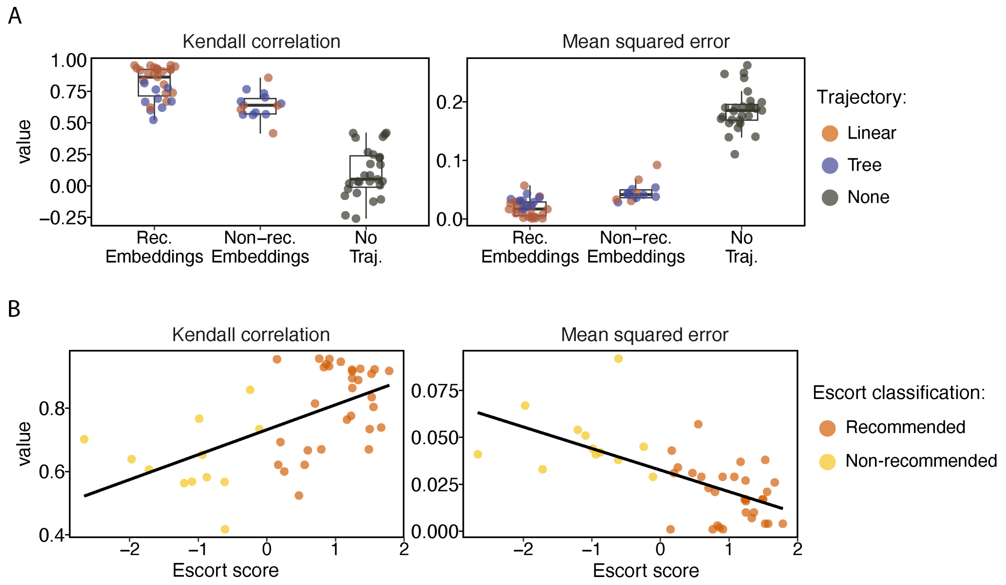


**Supplemental Figure 1. Assessment of Escort's performance on simulated datasets with trajectories generated by Monocle3. A.** The accuracy of trajectories generated by nine different embedding options was evaluated for eight simulated single-cell datasets. The trajectories were assessed using Kendall correlation and mean squared error. Different trajectory topologies are color-coded. The y-axis displays the values for the accuracy metric. **B**. Each embedding's Escort score (x-axis) versus the value for each accuracy metric (y-axis) is shown and colored according to their classification by Escort.

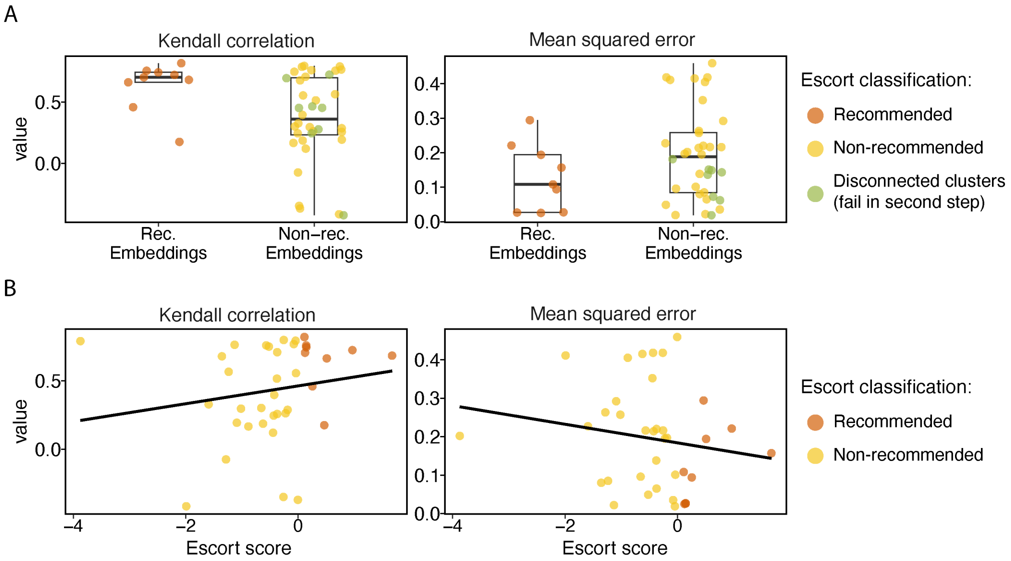


**Supplemental Figure 2. Assessment of Escort's performance on public scRNA-seq datasets with trajectories generated by Monocle3. A.** The accuracy of trajectories generated by nine different embedding options was evaluated for five publicly available scRNA-seq datasets. The trajectories were assessed using Kendall correlation and mean squared error. The colors distinguish each embedding classification by Escort, in addition to those embeddings that failed in the second step. The y-axis displays the values for the accuracy metrics. The x-axis corresponds to recommendations generated by Escort. **B**. Similar to A with the x-axis showing the Escort score.

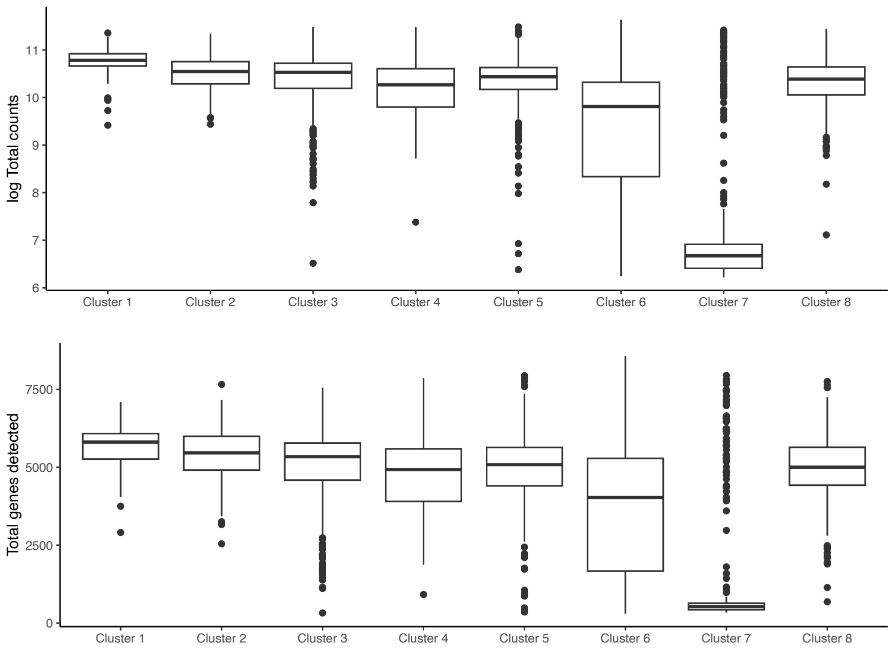


**Supplemental Figure 3.** Log of the total counts (top) and total genes detected per cell (bottom) across clusters in the hypertrophic chondrocyte scRNA-seq dataset. Cluster 7 was removed as these appeared to be low-quality.

**Supplementary Table 1.** Escort scores for all embeddings evaluated for the hypertrophic chondrocyte scRNA-seq dataset using either Slingshot (SS) or Monocle3 (M3) to build the preliminary trajectory. The original embedding option is highlighted in gray.

| Trajectory | Dimension reduction | # highly variable genes | Escort Score | Relative Ranking | Escort Decision |
|---|---|---|---|---|---|
| SS | PCA | 2006 (10% genes) | 0.1 | 1 | Recommended |
| SS | PCA | 4011 (20% genes) | 0.068 | 2 | Recommended |
| SS | PCA | 10028 (50% genes) | 0.009 | 3 | Recommended |
| SS | MDS | 2006 (10% genes) | -0.03 | 4 | Non-recommended |
| SS | MDS | 4011 (20% genes) | -0.098 | 5 | Non-recommended |
| SS | MDS | 10028 (50% genes) | -0.189 | 6 | Non-recommended |
| M3 | MDS | 10028 (50% genes) | -0.878 | 7 | Non-recommended |
| M3 | PCA | 4011 (20% genes) | -0.931 | 8 | Non-recommended |
| M3 | MDS | 4011 (20% genes) | -1.174 | 9 | Non-recommended |
| M3 | PCA | 2006 (10% genes) | -1.369 | 10 | Non-recommended |
| M3 | UMAP | 4011 (20% genes) | -1.488 | 11 | Non-recommended |
| M3 | MDS | 2006 (10% genes) | -1.547 | 12 | Non-recommended |
| M3 | PCA | 10028 (50% genes) | -1.634 | 13 | Non-recommended |
| M3 | UMAP | 10028 (50% genes) | -1.675 | 14 | Non-recommended |
| M3 | TSNE | 2006 (10% genes) | -1.725 | 15 | Non-recommended |
| M3 | TSNE | 4011 (20% genes) | -2.05 | 16 | Non-recommended |
| M3 | PCA+UMAP | 2000 | -2.082 | 17 | Non-recommended |
| M3 | TSNE | 10028 (50% genes) | -2.623 | 18 | Non-recommended |
| SS | TSNE | 4011 (20% genes) | -3.393 | 19 | Non-recommended |
| SS | PCA+UMAP | 2000 | -3.728 | 20 | Non-recommended |

| | | | | | |
|---|---|---|---|---|---|
| SS | TSNE | 10028 (50% genes) | -3.735 | 21 | Non-recommended |
| SS | UMAP | 4011 (20% genes) | -3.774 | 22 | Non-recommended |
| SS | UMAP | 10028 (50% genes) | -3.835 | 23 | Non-recommended |
| SS | TSNE | 2006 (10% genes) | -4.177 | 24 | Non-recommended |
| M3 | UMAP | 2006 (10% genes) | NA | NA | Non-recommended |
| SS | UMAP | 2006 (10% genes) | NA | NA | Non-recommended |
